# Genomic sequence capture of haemosporidian parasites: Methods and prospects for enhanced study of host-parasite evolution

**DOI:** 10.1101/420414

**Authors:** Lisa N. Barrow, Julie M. Allen, Xi Huang, Staffan Bensch, Christopher C. Witt

## Abstract

Avian malaria and related haemosporidians (*Plasmodium, [Para]Haemoproteus*, and *Leucocytoozoon*) represent an exciting multi-host, multi-parasite system in ecology and evolution. Global research in this field accelerated after 1) the publication in 2000 of PCR protocols to sequence a haemosporidian mitochondrial (mtDNA) barcode, and 2) the development in 2009 of an open-access database to document the geographic and host ranges of parasite mtDNA haplotypes. Isolating haemosporidian nuclear DNA from bird hosts, however, has been technically challenging, slowing the transition to genomic-scale sequencing techniques. We extend a recently-developed sequence capture method to obtain hundreds of haemosporidian nuclear loci from wild bird samples, which typically have low levels of infection, or parasitemia. We tested 51 infected birds from Peru and New Mexico and evaluated locus recovery in light of variation in parasitemia, divergence from reference sequences, and pooling strategies. Our method was successful for samples with parasitemia as low as ∼0.03% (3 of 10,000 blood cells infected) and mtDNA divergence as high as 15.9% (one *Leucocytozoon* sample), and using the most cost-effective pooling strategy tested. Phylogenetic relationships estimated with >300 nuclear loci were well resolved, providing substantial improvement over the mtDNA barcode. We provide protocols for sample preparation and sequence capture including custom probe kit sequences, and describe our bioinformatics pipeline using aTRAM 2.0, PHYLUCE, and custom Perl and Python scripts. This approach can be applied to the tens of thousands of avian samples that have already been screened for haemosporidians, and greatly improve our understanding of parasite speciation, biogeography, and evolutionary dynamics.

## Introduction

Multi-host, multi-parasite systems provide extensive opportunities to advance research in ecology and evolution. Haemosporidians (malaria and relatives, Order Haemosporida), the intracellular, protozoan parasites that infect vertebrates, are one great example, with studies ranging in scope from regional and temporal patterns of community turnover (e.g., Fallon *et al*. 2004, 2005; Olsson-Pons *et al*. 2015; Fecchio *et al*. 2017), to host-switching and diversification across long evolutionary timescales (e.g., Martinsen *et al*. 2008; Ricklefs *et al*. 2014; Galen *et al*. 2018a; Pacheco *et al*. 2018). Avian haemosporidians in particular (genera *Plasmodium*, [*Para*]*Haemoproteus*, and *Leucocytoozoon*) have attracted a large research community seeking to describe global patterns of diversity, abundance, and host range, and uncover mechanisms underlying parasite diversification, host-switching, and host susceptibility (e.g., Scheuerlein & Ricklefs 2004; Bensch *et al*. 2009; Clark *et al*. 2014; Lutz *et al*. 2015). The latter goal has particular importance for avian conservation, as exemplified by the Hawaiian honeycreepers, which have been severely impacted by the introduction of avian malaria (van Riper *et al*. 1986; Atkinson & LaPointe 2009).

The detection and description of avian blood parasites have accelerated with the application of molecular methods. While microscopy of thin blood smears remains essential for morphological verification and detailed species descriptions (Valkiunas 2005; Valkiunas *et al*. 2008), the field benefited substantially from the development of PCR primers for avian haemosporidians (Bensch *et al*. 2000; Fig. 1). Subsequent nested PCR protocols based on these primers (Hellgren *et al*. 2004; Waldenström *et al*. 2004) enable researchers to amplify and sequence a mitochondrial (mtDNA) barcode fragment, 478 base pairs of cytochrome *b* (*cytb*), from avian blood or tissue samples, even when infection levels (i.e., parasitemia) are too low to detect by microscopy. Parasite barcode sequences can then be compared with and uploaded to the avian haemosporidian database, MalAvi (Bensch *et al*. 2009). The growth of this database over the last decade has allowed for global analyses of parasite distributions and community assembly (Clark *et al*. 2014, 2017; Clark 2018; Ellis & Bensch 2018). It has become clear, however, that incorporating multiple nuclear loci will be necessary to further advance haemosporidian research.

**Fig. 1.**
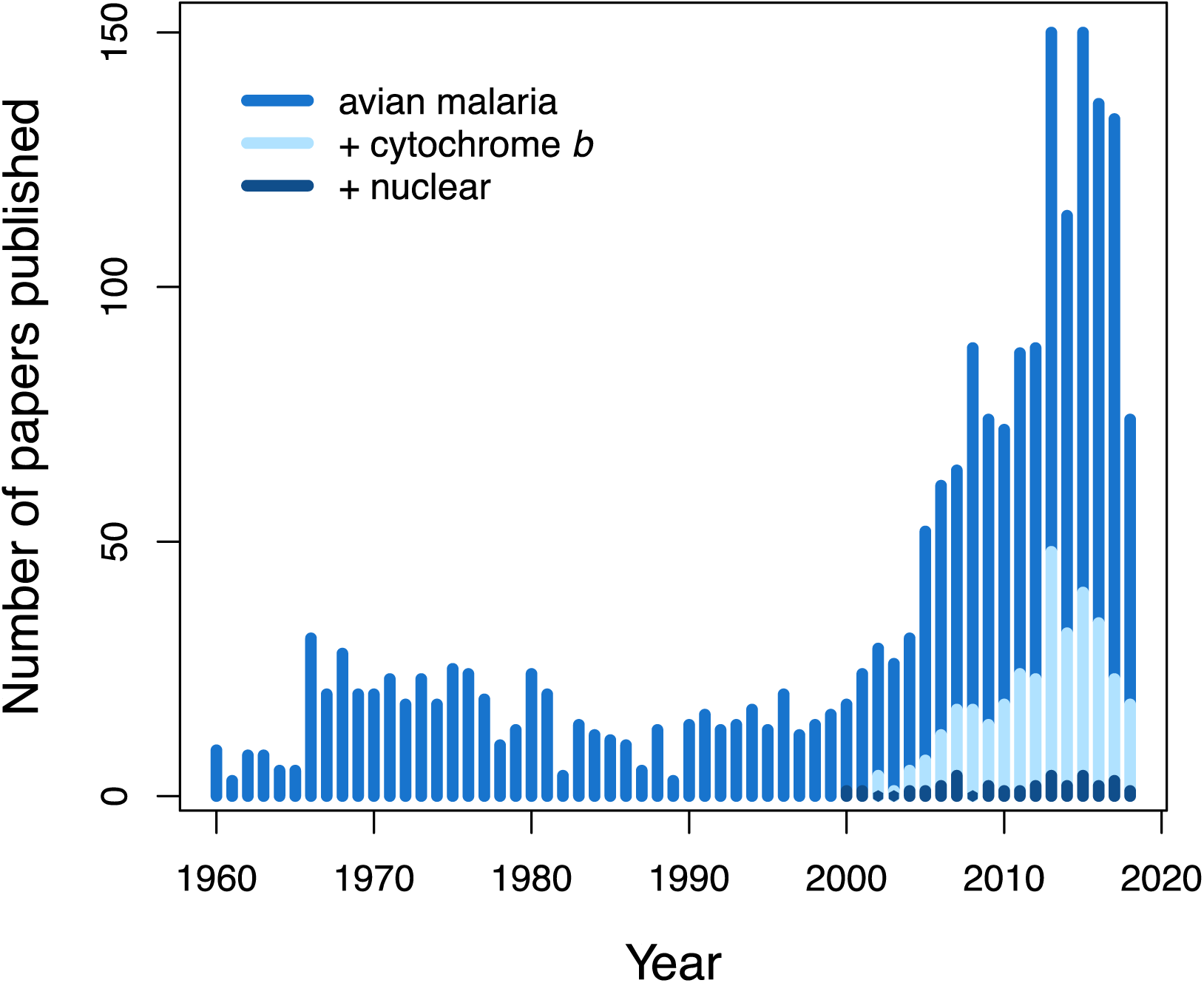
Papers published on ‘avian malaria’ by year since 1960 (n = 2,066) based on a Web of Science search (Clarivate Analytics, August 2018). Since 2000, many more ‘avian malaria’ papers include ‘mitochondrial’ (not shown, n = 218) or the barcode gene ‘cytochrome *b*’ (n = 339) than include ‘nuclear’ (n = 32).

Relatively few studies thus far have included multi-locus nuclear data of haemosporidian parasites (Fig. 1). Studies demonstrate that the *cytb* barcode provides limited resolution; a single *cytb* haplotype can include multiple cryptic species (Falk *et al*. 2015; Galen *et al*. 2018b), and phylogenies estimated from multiple nuclear loci substantially improve inferences of evolutionary relationships (Borner *et al*. 2016; Galen *et al*. 2018a). Several challenges, however, have previously prevented any large-scale efforts to obtain genomic data from avian haemosporidians. In contrast to mammalian red blood cells, avian red blood cells are nucleated, and the ratio of host to parasite DNA can be as high as a million to one (Perkins 2014). High-throughput sequencing of genomic DNA from avian samples is thus inefficient. Isolation of parasite gametocytes is possible through laser microdissection microscopy (Palinauskas *et al*. 2010), though this process is time-consuming and requires sufficient material. Parasite and host DNA can also be separated by inducing in vitro exflagellation of gametocytes, followed by centrifugation (Palinauskas *et al*. 2013), but donor birds with fairly high levels of infection are needed. Furthermore, avian haemosporidians are highly divergent from one another, with mtDNA barcode divergences between genera of ∼10–20%, making the task of designing general primers for multiple nuclear loci somewhat intractable. An ideal method would: 1) work on any sample that contained preserved DNA, including the tens of thousands of frozen blood and tissue samples that have been screened for haemosporidians, 2) be broadly applicable across a diverse group of haemosporidian parasites, and 3) enable cost-effective sequencing of hundreds of haemosporidian nuclear loci.

Sequence-capture methods used in conjunction with high-throughput sequencing are rapidly resolving the evolutionary Tree of Life for a variety of vertebrate, invertebrate, and plant taxa (Faircloth *et al*. 2012; Lemmon *et al*. 2012; Buddenhagen *et al*. 2016; Hamilton *et al*. 2016; Faircloth 2017; Quattrini *et al*. 2018). These techniques allow for the enrichment of genomic regions of interest by hybridizing oligonucleotide probes to genomic samples and removing non-target regions prior to sequencing (Albert *et al*. 2007; Gnirke *et al*. 2009). Probe sets can be designed from any existing genomic resources and are often useful across divergent taxa, although locus recovery tends to decline with increasing levels of divergence (Lemmon *et al*. 2012; Huang *et al*. 2018). The first genome for an avian *Haemoproteus* parasite was published in 2016 (Bensch *et al*. 2016), and no genomes for *Leucocytozoon* are available thus far. Given the vast differences between bird and haemosporidian genomes, the prospects are promising for targeted sequence capture of parasite genes from infected bird samples.

Huang *et al*. (2018) applied the first sequence-capture assay to haemosporidians, including eight *Haemoproteus* and one *Plasmodium* lineage, primarily from Europe and Asia. They successfully sequenced >100 nuclear exons from samples with up to ∼6% mtDNA divergence from the reference, *H. tartakovskyi*. It is not yet known, however, whether this approach will be useful for sequencing low-level infections that are most commonly observed in naturally infected birds. The minimum parasitemia tested in Huang *et al*. (2018) was 0.25%, while most naturally infected birds exhibit parasitemia less than 0.1% (Atkinson *et al*. 2001; Zehtindjiev *et al*. 2008; Ishtiaq *et al*. 2017). It is possible that when parasitemia is too low, parasite DNA will be overwhelmed by host bird DNA.

Our primary goal was to design a cost-effective sequence capture assay to work broadly across the genus *Haemoproteus* because of its global abundance, diversity, and variation in host specificity. Secondarily, we include promising results from a single *Leucocytozoon* sample, and generate nuclear sequences for this genus that can be incorporated into subsequent probe designs. Our specific objectives were to: 1) describe the relationship between parasitemia levels and sequence-capture success, 2) test how sequence-capture success is affected by percent divergence from the reference sequences used for probe design, and 3) compare strategies for pooling samples before capture to increase cost-effectiveness. To facilitate use of this method by scientists studying avian haemosporidians, we provide detailed laboratory protocols, including probe-kit sequences. We also describe our bioinformatics pipeline using aTRAM 2.0 (Allen *et al*. 2015, 2018) for locus assembly, and PHYLUCE (Faircloth 2015) and custom PERL and Python scripts for downstream processing.

## Materials and Methods

### Locus selection and probe design

The recently sequenced genome of *H. tartakovskyi* was used for initial probe design and sequence capture, targeting 1,000 genes (Bensch *et al*. 2016; Huang *et al*. 2018). We used the parasite-enriched sequences from three *Haemoproteus* species produced with this initial probe kit as references to design the probe kit used in this study. These three species provide representative variation across a large portion of the *Haemoproteus* phylogeny, with up to 6.5% sequence divergence in the mtDNA *cytb* barcode between them.

Paired reads obtained from *H. tartakovskyi* (lineage SISKIN1, sample ID 126/11c), *H. majoris* (PARUS1, 1ES86798), and *H. nucleocondensus* (GRW01, 512022) captures were trimmed, set as paired reads, and mapped to the initial 1,000 *H. tartakovskyi* genes in Geneious 8.1.9 (Biomatters Ltd). We selected 498 loci that were successfully captured and sequenced for at least one of the non-*tartakovskyi* samples, using a threshold of 3X coverage with no mismatches and alignment lengths of at least 200 base pairs (bp) as cutoffs for success. Most loci (471 out of 498) included three reference species, and the average alignment lengths were 1,251 bp (range: 232–8,319; total targeted bp: 622,788; Supplementary Table S1). Locus alignments of the three species were submitted to MYcroarray (now Arbor Biosciences, Ann Arbor, MI) for design and synthesis of a custom MYbaits kit with 19,973 biotinylated RNA probes and 2X tiling.

### Sample selection and quantification

We selected 51 bird samples for sequence capture; 50 with putative single infections of known *Haemoproteus* lineages and one with a mixed infection of *Leucocytozoon* and *Haemoproteus*. All samples consisted of pectoral muscle previously collected from wild birds in Peru or New Mexico, USA in accordance with approved animal care guidelines and permits. Samples were stored at -80°C in the Museum of Southwestern Biology Division of Genomic Resources at the University of New Mexico. Genomic DNA was extracted using an Omega Bio-tek EZNA Tissue DNA Kit following manufacturer protocols. For initial assessment of infection, three nested PCR protocols were used to maximize detection of all three parasite genera (Hellgren *et al*. 2004; Waldenström *et al*. 2004). Positive infections were identified by visualizing PCR products on an agarose gel, and haplotypes were assigned by sequencing the 478-base pair haemosporidian mtDNA barcode (*cytb*), as described in Marroquin-Flores *et al*. (2017). We chose samples infected with 14 *Haemoproteus* lineages (9 Peru, 5 New Mexico) and one *Leucocytozoon* lineage from Peru for subsequent quantification, capture, and sequencing (Table 1; Table S2).

**Table 1.** Sampling information for haemosporidian parasite lineages included in the study. All are *Haemoproteus* except one *Leucocytozoon* (*Leuc*). Additional information for each specimen is provided in Table S2.

| Parasite lineage | Study region | N samples | N host species | % mtDNA divergence from *H. tartakovskiyi* | Nearest reference | % mtDNA divergence from nearest reference |
| --- | --- | --- | --- | --- | --- | --- |
| G001 | Peru | 18 | 18 | 4.1 | *H. majoris* | 3.77 |
| G002 | Peru | 5 | 3 | 5.57 | *H. tartakovskiyi* | 5.57 |
| G003 | Peru | 2 | 2 | 3.64 | *H. tartakovskiyi* | 3.64 |
| G004 | Peru | 2 | 2 | 6.2 | *H. majoris* | 5.23 |
| G005 | Peru | 2 | 2 | 2.57 | *H. tartakovskiyi* | 2.57 |
| G006 | Peru | 2 | 2 | 3.43 | *H. tartakovskiyi* | 3.43 |
| G007 | Peru | 6 | 5 | 3.21 | *H. tartakovskiyi* | 3.21 |
| G009 | Peru | 4 | 4 | 3.43 | *H. tartakovskiyi* | 3.43 |
| G011 | Peru | 3 | 3 | 3.0 | *H. tartakovskiyi* | 3.0 |
| G403 (*Leuc*) | Peru | 1 | 1 | 18.0 | *H. nucleocondensus* | 15.9 |
| CHOGRA01 | New Mexico | 1 | 1 | 1.71 | *H. tartakovskiyi* | 1.71 |
| SPIPAS01 | New Mexico | 2 | 2 | 1.71 | *H. tartakovskiyi* | 1.71 |
| SPISAL01 | New Mexico | 1 | 1 | 6.0 | *H. nucleocondensus* | 4.81 |
| TROAED12 | New Mexico | 1 | 1 | 5.9 | *H. majoris* | 3.77 |
| VIGIL05 | New Mexico | 1 | 1 | 4.5 | *H. majoris* | 3.98 |

Overall sample DNA concentrations were quantified using a Qubit 3.0 Fluorometer. Relative parasite DNA concentrations were assessed using quantitative PCR with primers targeting a 154-base pair portion of haemosporidian ribosomal RNA (Fallon *et al*. 2003). We used 2X iTaq Universal SYBR Green Supermix (Bio-Rad), 0.5 µM of each primer (343F and 496R), and 30 ng sample DNA in a total reaction volume of 20 µL. Reactions were run on a Bio-Rad CFX96 Real-Time PCR System with the following temperature profile: 95 °C for 3 min, 40 cycles of 95 °C for 15 sec and 57°C for 1 min, followed by a melt curve analysis (47 °C to 95 °C at 0.5 °C and 5 sec per cycle). Each plate included three no-template controls. To generate a standard curve, we made a six-step 1:10 serial dilution (30–0.0003 ng DNA per well) of one sample with high parasitemia as estimated by microscopy (NK168012; 1.86% cells infected). Each sample was run in triplicate and cycle threshold (CT) values were averaged across the three replicates.

### Library preparation, capture, and sequencing

We prepared libraries for each sample using the KAPA Hyper Prep Kit (Kapa Biosystems) and dual-indexing with the iTru system (Faircloth & Glenn 2012; Glenn *et al*. 2016; baddna.uga.edu). Complete protocols are provided in Supporting Information. Briefly, genomic DNA was visualized on an agarose gel to verify high molecular weight and determine a suitable sonication protocol. Samples were then fragmented to a length distribution centered on ∼500 bp using a Covaris M220 Focused-ultrasonicator (Covaris, Inc.). Libraries were prepared primarily following the KAPA Hyper Prep Kit protocols, using 250 ng of fragmented DNA per sample, custom indexed adapters, KAPA Pure Beads for bead clean-ups, and 10 cycles in the indexing amplification step. After quantifying libraries by Qubit, equal amounts of sample libraries were combined in pools of eight, four, two, or a single sample for capture. We only pooled libraries with similar parasitemia values as determined by qPCR in an attempt to obtain even capture success and sequencing coverage across samples within a pool. Capture pools contained 1–2 µg DNA (at least 125 ng per library).

Hybrid enrichment was performed following the MYbaits Version 3.02 protocols with minor modifications as follows. Block Mix 3 was prepared from custom oligos for the iTru dual-indexing system, and Chicken Hybloc DNA (Applied Genetics Laboratories, Inc.) was used as Block Mix 1. We extended the hybridization time to 36–40 hr, as recommended to increase capture efficiency for low-abundance targets. For the post-capture amplification, we used 2X KAPA HiFi HotStart ReadyMix with the bead-bound library and the following thermal profile: 98 °C for 2 min, 16 cycles of 98 °C for 20 sec, 60 °C for 30 sec, and 72 °C for 60 sec, followed by a final extension at 72 °C for 5 min and held at 4 °C. Post amplification, we removed the beads and performed a final 1.2X KAPA Pure Bead clean-up. Captured pools were quantified and characterized by Qubit and an Agilent 2100 Bioanalyzer, and shipped to the Oklahoma Medical Research Foundation (OMRF) Clinical Genomics Center for final qPCR and sequencing. All capture pools were combined and run on a single lane of PE150 Illumina HiSeq 3000.

### Bioinformatic processing

Demultiplexed reads for each sample were obtained from the OMRF Clinical Genomics Center. We trimmed adapters and low-quality bases using default settings in Illumiprocessor 2.0.6 (Faircloth 2013), which provides a wrapper for Trimmomatic (Bolger *et al*. 2014). These settings include trimming reads with lengths <40 (MINLEN:40), bases at the start of a read with quality scores <5 (LEADING:5), bases at the end of a read with scores <15 (TRAILING:15), and bases in a sliding window where four consecutive bases have an average quality <15 (SLIDINGWINDOW:4:15). We then used the automated Target Restricted Assembly Method, aTRAM 2.0 (Allen *et al*. 2015, 2018; http://www.github.com/juliema/aTRAM) to assemble haemosporidian parasite genes using the 498 reference *Haemoproteus* gene sequences. This approach uses local BLAST searches and an iterative approach to produce assemblies for genes of interest from cleaned read data. We used BLAST 2.7.1 (Altschul *et al*. 1990), Trinity 2.0.6 (Grabherr *et al*. 2011) as the assembler, five iterations, and nucleotide reference sequences from *H. tartakovskyi* for all individuals. We also conducted two additional tests to improve locus recovery for individuals with higher divergences from the reference. First, we used amino acid reference sequences instead of nucleotide for aTRAM assemblies, but found that several bird host genes were assembled in place of the haemosporidian genes. Second, we used *H. majoris* as the nucleotide reference, and added the new locus assemblies recovered to the set for further processing.

We next used custom scripts written in Perl and Python (available at https://github.com/juliema/) to keep only the contigs from the last aTRAM iteration, compare and align them to the translated exon sequences for the reference *H. tartakovskyi* using Exonerate 2.2.0 (Slater & Birney 2005), and stitch together any exons that were broken into multiple contigs (as described in Allen *et al*. 2017). Because aTRAM performs iterative assemblies that can extend outward from the original reference sequence, the last iteration is most likely to have the most complete, longest contigs. For samples where the middle of a locus was either not sequenced or not recovered due to low coverage, we used exon stitching to retain both ends of the sequence for alignment. We then conducted a reciprocal best-BLAST check on the stitched exons, and removed any individual-locus combination for which the top match for the assembled locus was not the target locus. For this search, we created a local BLAST database from all 6,436 *H. tartakovskyi* genes (downloaded from http://mbio-serv2.mbioekol.lu.se/Malavi/Downloads). Fewer than 0.2% (9 of 4,507) individual-locus combinations were mismatched and therefore removed. Prior to multiple sequence alignment, we added sequences for the three reference species, and used custom Python scripts (available on Dryad) to reformat the aTRAM sequences for the PHYLUCE pipeline (Faircloth 2015).

Several PHYLUCE scripts were used to summarize locus information for each individual, produce multiple sequence alignments, and generate concatenated datasets for phylogenetic analysis in RAxML. We considered samples with at least 50 recovered loci to be successful, and generated alignments including only those individuals. We generated edge-trimmed alignments with MAFFT 7.130b (Katoh & Standley 2013), using a threshold parameter of 0.3 (at least 30% of individuals with sequence at the edges) and maximum divergence of 0.4. As one final check, we manually examined all alignments with at least 50% of taxa (404 alignments), and removed 25 that were poorly-aligned. Concatenated RAxML alignments were generated allowing different levels of missing data: at least 50, 70, or 90% of individuals per locus.

### Sequence coverage estimation

To provide an estimate of sequence coverage and its influence on capture success, we used the BLAST-only option in aTRAM to find and count the reads for a parasite mtDNA gene and a host mtDNA gene for each sample. The *H. tartakovskyi* reference sequence was used to estimate coverage for the parasite mtDNA *cytb* gene. Given the availability of host bird sequences on GenBank, we used *ND2* reference sequences (Table S2) to estimate host mtDNA coverage. Host genes were not targeted by the probe kit, but these non-target reads were sequenced as a by-product because of the large quantity of bird mtDNA in the samples. We estimated per-site coverage based on the length of the reference gene used, assuming 150 base pair read lengths, and compared the ratio of parasite to host per-site coverage between samples that were considered capture successes and failures.

### Detection of mixed infections

For each successfully-captured individual, we mapped the cleaned, paired reads to both *COI* and *cytb* mtDNA reference sequences in Geneious to check for possible co-infections. In cases where multiple haplotypes were apparent in the mapped read assembly, we compared the reads to the *cytb* barcode region for the original assigned haplotype to sort out the alternative haplotype and determine the variant frequency.

### Downstream analysis

To test for effects of multiple variables on sequence capture success, we used generalized linear models (GLMs) in R (R Core Team 2016). We tested whether parasitemia (qPCR CT value), level of divergence from the nearest reference (% mtDNA sequence divergence), or the number of samples pooled (factor with two categories: 8 or <8), had an effect on the number of loci recovered per sample. We also included the interaction between parasitemia and divergence from the reference. The one *Leucocytozoon* sample was an outlier for mtDNA divergence, thus we repeated analyses with and without this individual.

To determine whether the nuclear loci we recovered improve inferences of haemosporidian relationships, we estimated a phylogeny for the samples with sufficient data. We used PartitionFinder 2 (Lanfear *et al*. 2017) to select appropriate models of nucleotide evolution and partitioning schemes for each dataset. Given the number of loci, we used the rcluster algorithm with RAxML (Stamatakis 2014; Lanfear *et al*. 2014), linked branch lengths, and AICc for model selection. Phylogenies were estimated for each nuclear dataset (50, 70, 90% complete matrices), for the mtDNA capture data (3,226 bp of COI and *cytb*), and for the original *cytb* barcode data (478 bp). We used the rapid hill-climbing algorithm in RAxML with the GTR+G model and 1000 bootstrap replicates. We also estimated species trees from the nuclear datasets using SVDquartets (Chifman & Kubatko 2014), implemented in PAUP* 4.0a163 (Swofford 2002). We performed an exhaustive search of all quartets and conducted multilocus bootstrapping with 1000 replicates and partitioned loci.

## Results

### Data summary

We obtained 620,951,640 reads from one sequencing lane, of which 571,186,910 (92%) were sorted by individual barcode. On average, 11.2 million (s.d.: ± 7.2 million) reads were obtained per individual (min–max: 3.7–39.3 million; median: 8.7 million). The number of parasite loci assembled per individual ranged from 491 (99%) to none; eight of the 51 samples resulted in no *Haemoproteus* loci. We considered 15 samples (29%) to be sequence capture successes, with >70 loci obtained. The remaining individuals resulted in 11 or fewer loci. For most of these failures, only one or two mtDNA genes were assembled. For the successful samples, an average of 295 ± 140 (min–max: 71–491, median: 254) loci were recovered with mean locus lengths of 2,428 (min–max: 2,317–2,798) bp. The average number of loci assembled with >1,000 bp in length was 241 ± 108 (min–max: 64–386; median: 212; Fig. S1).

### Effects of parasitemia, divergence, and pooling on success

Parasitemia was positively correlated with capture success and locus recovery (t = 6.15, p < 0.0001; Fig. 2a). Samples with parasitemia values >0.07% (∼7 out of 10,000 infected cells) were all successful, and samples with as low as ∼0.03% (∼3 of 10,000 cells; qPCR CT value ∼26) also resulted in >100 loci. Divergence from the nearest reference was negatively correlated with locus recovery (t = -3.91, p = 0.0003; Fig. 2b). There was also a significant interaction between parasitemia and divergence (t = -3.76, p = 0.0005). Samples with both sufficient parasitemia and low divergence from the reference had the best locus recovery (>400 loci). The most divergent capture success was the *Leucocytozoon* sample, with 15.9% mtDNA divergence from the nearest reference and 71 loci recovered. GLM results excluding this outlier were qualitatively similar but stronger in magnitude. The number of samples included in a pool did not affect capture success (t = 0.68, p = 0.5); several individuals in the most cost-effective, 8-sample pools resulted in >100 sequenced loci.

**Fig. 2.**
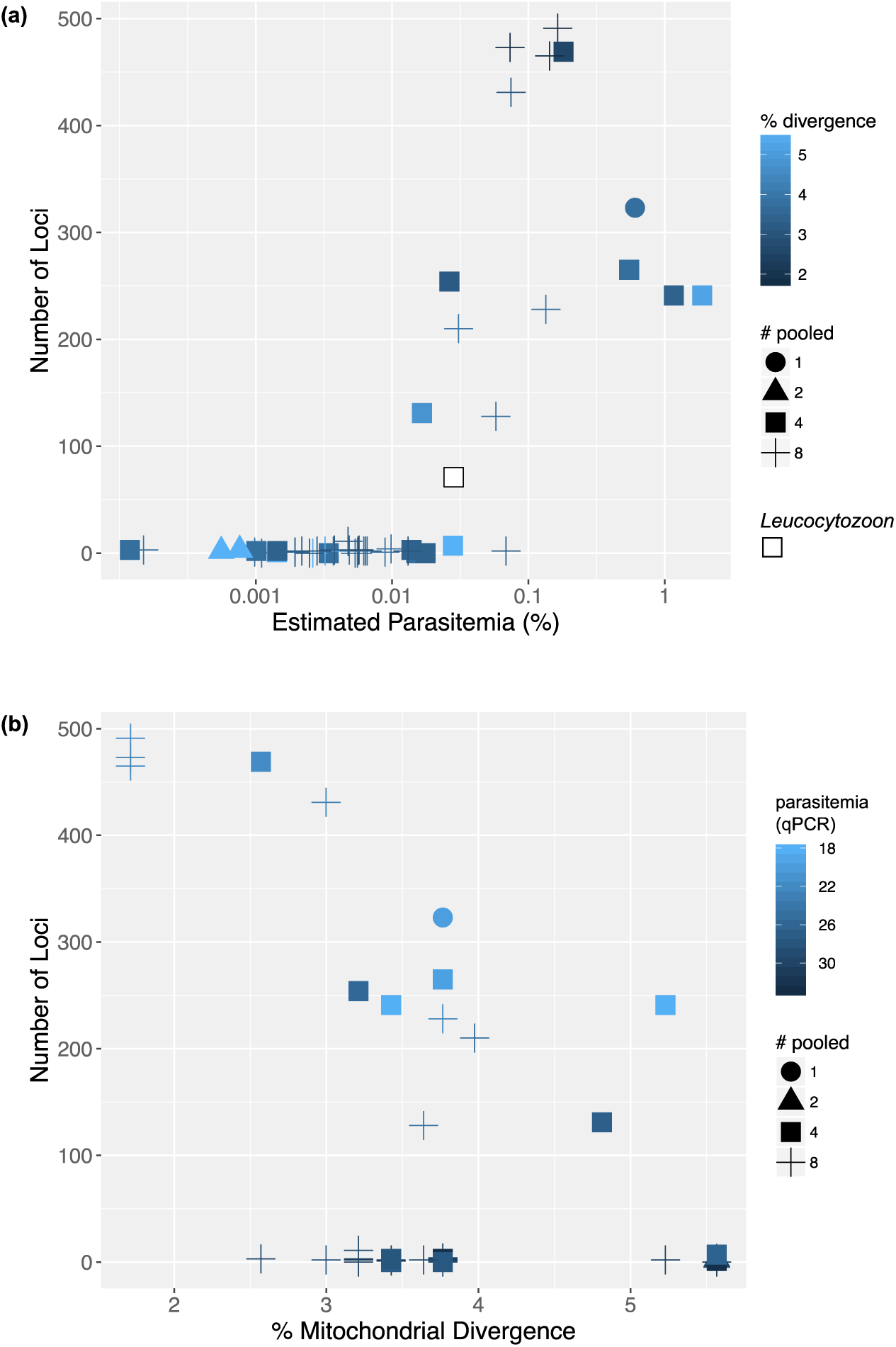
Number of parasite loci sequenced out of 498 total. (a) Locus recovery increased with parasitemia, shown as % infected cells estimated by microscopic examination of the qPCR standard. Colors depict % divergence from the nearest reference. (b) Locus recovery decreased with increasing % divergence from references, but with sufficient parasitemia, divergent samples were successful. Colors depict parasitemia as the qPCR CT value. The *Leucocytozoon* sample (15.9% divergent) is not shown in (b) to improve visualization. In both plots, shapes depict the number of samples in a capture pool, which did not influence locus recovery.

### Parasite versus host coverage

The estimated depth of sequence coverage per site for parasite mtDNA showed substantial variation across samples, with a mean of 2,804 reads (min–max: 0.39–45,683). Samples that were considered capture successes had a mean parasite mtDNA per-site coverage of 9,516 (185– 45,683) and mean host mtDNA coverage of 856 (1.73–2,540). In contrast, samples that were considered capture failures had a mean parasite mtDNA coverage of only 8.22 (0.39–125), while host mtDNA coverage was similar with a mean of 957 (0.87–3,110). On average, the ratio of parasite to host coverage was 137 (0.09–1,622) for capture successes, and only 0.32 (0.0002– 11.15) for capture failures.

### Detection of mixed infections

We identified three samples co-infected with multiple lineages, two of which had not been previously detected by nested PCR and Sanger sequencing. Sample NK168883 was co-infected with the *Haemoproteus* lineage G009 and a novel *Leucocytozoon* lineage (assigned name G403). Within the MalAvi barcode region, the frequency of the *Leucocytozoon* lineage dominated the reads and ranged from 94.8–98.0%. The concatenated sequence for this sample had >15% sequence divergence from the pure G009 *Haemoproteus* sample (NK168881), indicating that the genes assembled for that sample belonged to *Leucocytozoon*. Sample NK275890 was co-infected with two *Haemoproteus* lineages, SPIPAS01 and SIAMEX01 (the read frequency of SIAMEX01 was 11.0–14.2%; mtDNA divergence between the two haplotypes of 6.1%). Sample NK276102 was also co-infected with two *Haemoproteus* lineages, VIGIL05 and VIGIL07 (the read frequency of VIGIL07 was 30.3–40.5%; mtDNA divergence 3.6%). Phylogenetic analyses were repeated without the two *Haemoproteus* co-infected samples because the loci extracted by aTRAM may have represented a composite of the two haplotypes that could not be definitively sorted.

### Phylogenetic resolution

The nuclear datasets substantially improved phylogenetic resolution of *Haemoproteus* parasites (Fig. 3). Most relationships were consistent among datasets with different levels of completeness. The 50% complete matrix included 377 loci and 287,164 bp (Fig. 3a), the 70% matrix included 206 loci and 189,883 bp, and the 90% matrix included 59 loci and 70,611 bp (Fig. 3b). The topologies resulting from RAxML and SVDquartets were identical for the 50% and 70% datasets. The position of *H. tartakovskyi* differed for the 90% matrix RAxML analysis (Fig. 3b), while the SVDquartets topology was consistent with the other datasets. Phylogenetic analyses excluding the *Haemoproteus* mixed infections resulted in similar topologies, except for the uncertain position of TROAED12 (Fig. S2). The majority of nodes had high support for the nuclear datasets; bootstrap values were ≥ 95 for 15 (94%) nodes with the 50% matrix and 13 (81%) nodes with the 90% matrix. In contrast, the two-gene mtDNA dataset and the *cytb* barcode produced poorly-resolved phylogenies for the lineages in our study, inferring very few relationships with any certainty (Fig. 3c,d). Only 6 (38%) and 3 (19%) of the nodes had bootstrap values ≥ 95, respectively.

**Fig. 3.**
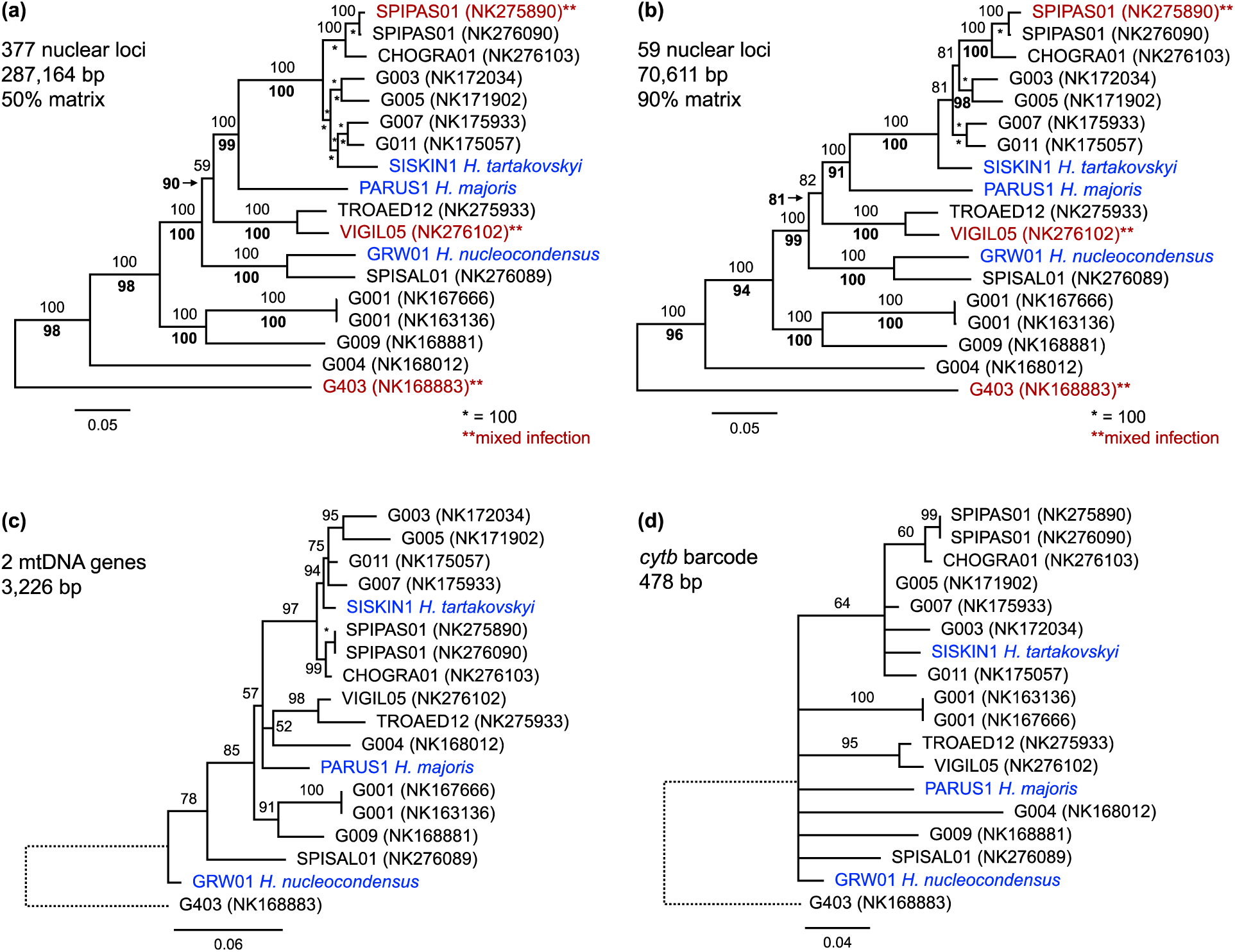
Phylogenies estimated with nuclear (a, b) and mitochondrial (c, d) datasets. Blue tip labels indicate reference samples. RAxML best trees are shown with branch lengths in substitutions per site and RAxML bootstrap support above branches. (a, b) SVDquartets support values are shown in bold below branches. Red tip labels indicate samples with mixed infections. (c, d) Branches with support values <50 were collapsed. Branch lengths for G403 (*Leucocytozoon*) were reduced to improve visualization.

## Discussion

### Capture success and parasitemia

The extremely low abundance of parasite DNA compared to host DNA has presented a great challenge for obtaining haemosporidian genomic data from naturally-infected birds. We provide parameters for the successful implementation of a new sequence-capture assay to obtain hundreds of haemosporidian parasite loci from wild bird samples. By quantifying parasitemia using either a standard qPCR protocol or microscopic examination, researchers can select samples above a certain threshold to enable capture success. Based on our results, samples with ∼0.07% parasitemia were always successful, and samples with as low as ∼0.03% parasitemia also tended to be successful.

This finding is exciting because tens of thousands of avian blood and tissue samples exist in museums and laboratories globally and have already been screened for haemosporidians. Our results indicate that a large portion of these samples (29% in our study) will be suitable for sequence capture of haemosporidian parasites. One possible improvement for sequence capture of samples with even lower parasitemia is to perform a double enrichment using the haemosporidian probe kit, in order to increase the relative amount of parasite DNA in the samples further. If a single sample is captured at a time, parasite and host DNA could accurately be quantified for each sample with qPCR before and after each enrichment. For a more cost-effective approach, however, we recommend quantifying parasitemia with qPCR prior to capture, and pooling sample libraries with similar values as we have done here, in order to obtain more even sequencing coverage across samples.

### Potential improvements and cost

Our probe kit was designed to work broadly across *Haemoproteus* because of our interest in the diversity and host range variation of this genus worldwide (Clark *et al*. 2014; Ellis & Bensch 2018) and the genomic resources available to design probes (Bensch *et al*. 2016; Huang *et al*. 2018). A clear extension of this method is to incorporate probes targeting all avian haemosporidian genera. Although we have not yet tested our probe kit on *Plasmodium*-infected birds, the successful capture and sequencing of >70 loci for *Leucocytozoon* is quite promising because avian *Plasmodium* is less divergent from *Haemoproteus*. Additional tests on *Leucocytozoon*-and *Plasmodium*-infected samples with the current probe kit can be carried out to determine success rates, and the generated sequences can be incorporated into new probe designs along with new and existing transcriptome and genome data for the other genera (Lutz *et al*. 2016; Videvall *et al*. 2017; Böhme *et al*. 2018).

As reference data for more parasite lineages are generated, it will likely be possible to improve certain steps of our bioinformatics pipeline. We tested two different reference species for aTRAM assemblies, and found that the reference chosen for assembly has some effect on the number of loci recovered; initial locus recovery was better for samples that were less divergent from the *H. tartakovskyi* reference. We assembled >425 loci (>85%) for the samples with the lowest divergence from *H. tartakovskyi* (1.7–3% mtDNA divergence), even though they did not have the highest parasitemia values. For the samples with higher parasitemia but higher divergences from *H. tartakovskyi* (up to 6.2% for *Haemoproteus* sample), we still recovered more than 200 loci. We were also able to add 45–60 more loci for these divergent samples by using *H. majoris* as a reference. In aTRAM, protein sequences can be used instead of nucleotide sequences as references for assembling more divergent loci, but in our case, contamination from host bird DNA resulted in some gene assemblies for the bird instead of the parasite. One potential work-around may be to filter reads by GC content prior to assembly, because avian haemosporidian genomes have lower GC content on average than bird hosts (Galen *et al*. 2018a), but this potential approach will require further testing.

One other challenge for haemosporidian research is that mixed infections are extremely common in nature. We could be fairly confident that the sequences from the co-infected *Leucocytozoon* sample did indeed belong to *Leucocytozoon* because we were able to compare them with a pure infection of the same *Haemoproteus* lineage. The two other co-infected samples in our study were not unambiguously sorted, but with reference sequences from one or both of the lineages in a mixed infection, a step could be added to the pipeline to sort them bioinformatically. One other future improvement to the pipeline will be to incorporate protein-guided multiple sequence alignments. We chose to manually check our MAFFT alignments and we removed 25 genes (6%) that were poorly-aligned. Because we are targeting exons, protein-guided DNA alignments may improve results for more divergent sequences and remove the need for extensive manual checking.

The total cost for our study including DNA extraction, qPCR, library preparation, capture, and sequencing was approximately $87 USD per sample, or $0.30 per locus based on the average of 295 loci per successful sample. For just library preparation and sequence capture, with eight libraries pooled before capture, we spent ∼$35 per sample, or $0.12 per locus. Costs could be reduced further by ordering larger capture kits or combining more samples together on one higher-output sequencing platform.

### Future directions for avian haemosporidian research

Avian haemosporidian research has grown at a rapid pace since molecular tools have been applied to the field. To date, more than 3,100 parasite mtDNA haplotypes have been discovered and uploaded to the MalAvi database along with their associated locality and host species. Broad syntheses of these data have provided important insights into global distribution patterns and host-parasite associations (Clark *et al*. 2014, 2017; Ellis & Bensch 2018), but the improved resolution of *Haemoproteus* relationships afforded by our genomic sequence capture data has great potential for moving avian haemosporidian research forward. First, the species limits of haemosporidian parasites are difficult to define. Some lineages differing by a single nucleotide in the *cytb* barcode region are considered to be reproductively isolated, biological species (Nilsson *et al*. 2016), while others are considered to represent intraspecific variants (Outlaw & Ricklefs 2014; Hellgren *et al*. 2015). Galen *et al*. (2018b) used seven nuclear loci to show that both phenomena occur in *Leucocytozoon*, confirming the poor resolution of mtDNA for inferring species limits. Second, developing large, multi-locus nuclear DNA sequence datasets is needed to advance the study of haemosporidian evolutionary dynamics. Robust phylogenies will allow for more accurate estimates of biogeographic history (Hellgren *et al*. 2015), trait evolution (Ellis & Bensch 2018), and transitions between host-generalist and host-specialist strategies (Loiseau *et al*. 2012b). In this way, methods for collecting haemosporidian genomic data will facilitate detailed studies of parasite diversification, host breadth, and distributional limits across the globe. Furthermore, high-resolution determination of haemosporidian species limits will be critical to identify and manage the novel host-parasite interactions that are expected to pose a major threat to host-species persistence during climate warming (Garamszegi 2011; Loiseau *et al*. 2012a).

## Acknowledgements

This work was supported by NSF DEB-1146491, a New Mexico Ornithological Society research grant, and an NSF Postdoctoral Fellowship (NSF PRFB-1611710) to LNB. We thank Michael Andersen, Jenna McCullough, and the staff of the UNM Center for Advanced Research Computing.

## Data Accessibility

Specimen information is available from the Arctos database (arctosdb.org). Parasite sequences are available on MalAvi and GenBank (Accession Numbers in Table S2). Protocols and probe sequences are included as supporting information. Scripts, sequence data, and alignments will be made available on Dryad.

## Author Contributions

LNB designed the study, collected the data, and led the writing; LNB and JMA analyzed the data; XH, SB, and CCW contributed genomic data and resources; all authors edited the manuscript.

